# Interpretation of network-based integration from multi-omics longitudinal data

**DOI:** 10.1101/2020.11.02.365593

**Authors:** Antoine Bodein, Marie-Pier Scott-Boyer, Olivier Perin, Kim-Anh Lê Cao, Arnaud Droit

## Abstract

Cost reduction of high-throughput technologies has enabled the monitoring of the same biological sample across multiple omics studies and multiple timepoints. The goal is to combine longitudinal multi-omics data to detect temporal relationships between molecules and interactions between omics layers. This can finally lead to uncover new regulation mechanisms and interactions that could be responsible for causing complex phenotype or disease. However multi-omics integration of diverse omics data is still challenging due to heterogeneous data and designs. Moreover, interpretation of multi-omics models is the key to understand biological systems.

We propose a generic analytic and integration framework for multi-omics longitudinal datasets that consists of multi-omics kinetic clustering and multi-layer network-based analysis. This framework was successfully applied to two case studies with different experimental designs and omics data collected. The first case studied transcriptomic and proteomic changes during cell cycle in human HeLa cells, while the second focused on maize transcriptomic and metabolomic response to aphid feeding. Propagation analysis on multi-layer networks identifies regulatory mechanisms and function prediction for both case studies.

Our framework has led to the identification of new multi-layer interactions involved in key biological functions that cannot be revealed with single omics analysis and interplay in the kinetics that could help identify novel biological mechanisms.

## 1 Introduction

Cost reductions of DNA sequencing in addition to other high-throughput multi-omics technologies have revolutionized many research fields ranging from personalized medicine [6, 12] to systems biology [10, 45]. These innovations have led to new biological insights and a better understanding of living organisms [15, 25, 41]. Thus, enabling the assessment of most biological layers, this democratisation of high-throughput technologies has created large datasets representing different biomolecules that necessitate specific processing and statistical methods [14, 32]. Multi-omics trials typically collect different types of biomolecules (mRNA, proteins, metabolites, etc.) from the same biological samples with the ultimate goal of highlighting the interaction between biological layers that could be responsible for causing complex phenotype or diseases [12]. Even more so most biological phenomenon involves complex interactions between layers that vary through time [22]. Adapted multi-omics time-course methods to integrate and accurately capture interactions among those biological layers are thus now required of and fully capture interactions within and between omic layers.

To describe complex interactions and regulatory mechanisms behind biological systems, mathematical models, such as network, are built to interpret and reverse-engineer cellular functions. Networks are used to represent all relevant interactions taking place in a biological systems [2]. In networks, molecules (genes, proteins, metabolites) are reduced to a series of nodes that are connected to each other by edges. Edges represent the pairwise relationships, interactions, between two molecules within the same network. Molecular networks have become extremely popular and have been used in every area of biology to model for example transcriptional regulation mechanisms, physical protein-protein interactions [17, 39], or metabolic reactions [18]. Networks come with valuable properties and useful topological features such as degree distribution to identify highly connected nodes or shortest paths which determine proximity between two nodes. On a different scale, network modularity defines sub-network units with highly connected nodes in respect to the rest of the network. These sub-networks, also known as modules, often share a similar function. Thus, the “guilt by association” property assumes that known or unknown highly connected molecules should be functionally related [54].

Inference methods for network construction are often applied to a single omic layer to identify interaction between molecules. However, this does not directly elucidate interaction across multiple omic layers [49]. To connect these layers, a first approach require prior knowledge of across omic molecular networks such as publicly-available databases [20]. This approach is based on a legacy limited to model organisms and may not reflect the current biological condition. A second method use multivariate data-driven methods that statistically infers correlations between molecules based on multi-omics data. However, this approach may have many possible solutions. A combination of the two methods could improve multi-omics network construction.

The ultimate goal of multi-omics networks is to connect phenotypes to biological mechanisms and their regulators. Analysis of the interactome identifies direct neighbors and modules linked to a phenotype. However, our knowledge of the interactome is not complete and direct connections can lead to false positive discoveries. Recently, propagation algorithms have become the state-of-the-art to investigate gene disease associations and also gene function prediction [8, 30]. Based on the known association, the signal is iteratively propagated through the network. When a steady state is reached, new nodes can be added to the initial association by their propagation score reflecting their proximity to the starting nodes. It thus highlights potential new phenotype-related targets. New advances in randoms walks algorithms allow to propagate the signal in heterogeneous multi-layered networks which improves association prediction [44].

In this paper, we propose to build hybrid multi-omics networks from longitudinal multi-omics data in order to facilitate the interpretation of multi-layers systems. (Figure 1). This methodology is based in the first place on the modelling and clustering of expression profiles with similar behaviours over time. It relies on both accurate network reconstruction methods and knowledge-based reviewed interactions between either molecules of the same or different types. Finally, a random walk algorithm was used to identify and make new hypothesis about links between omics molecules and key biological functions or mechanisms. We illustrate this approach through 2 case studies. These studies have different experimental designs with different omics data types, timepoints and organisms to demonstrate that the proposed approach is able to deal with a wide range of situations.

**Figure 1:**
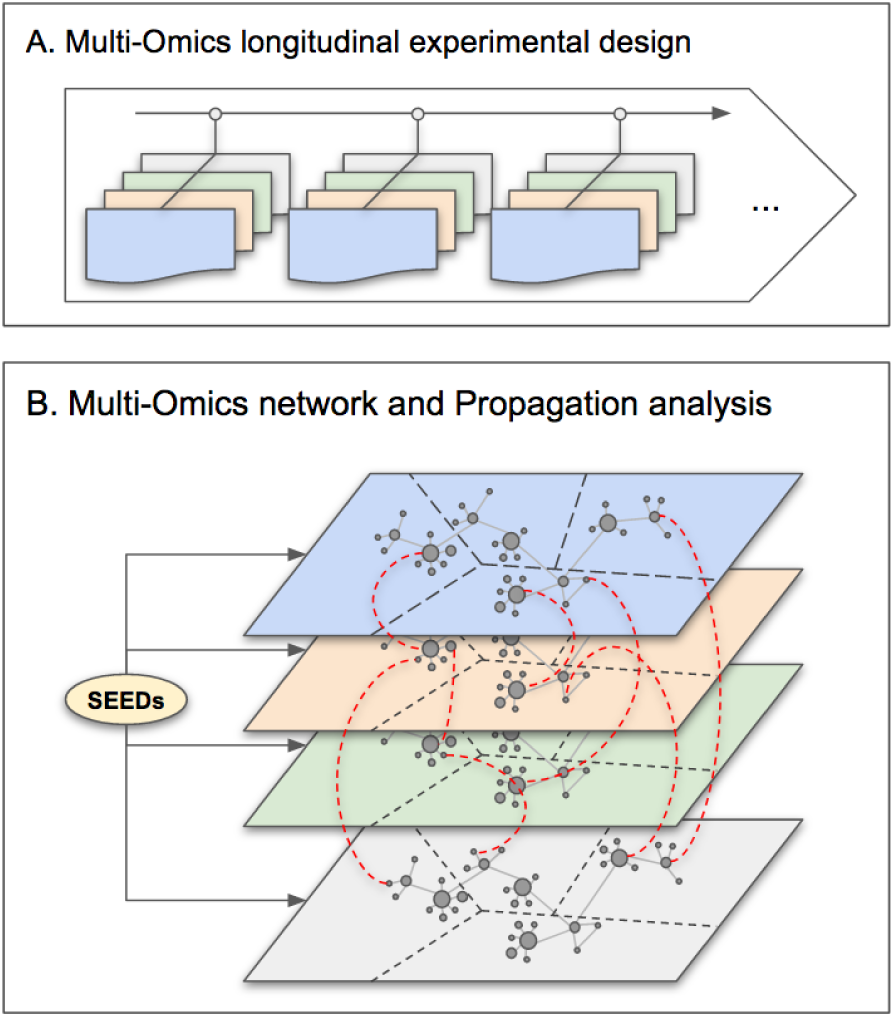
Overview of the proposed approach. **(A)** Description of the experimental design: the same biological material is sampled at several time points across several omic layers indicated in different colors. Each omic data is normalised using both platform-specific and time-specific normalisation steps. **(B)** Multi-Omics Network is built using both inference-based and knowledge-based methods to connect intra- and cross-layered biological features or molecules (mRNA, proteins, metabolites). Measured molecules are clustered into groups of similar expression profiles over time and corresponding nodes formed kinetic sub-networks. Over-representation analysis is performed to add an extra layer of functional annotation. Propagation analysis is performed on specific nodes of interest, called *seeds* (biological function, gene, protein, metabolite, etc.) to identify closely related molecules.

## 2 Material and Methods

This approach proposes pre-processing, modelling and clustering steps for multi-omics longitudinal data. It mainly emphasizes on network-based integration and multi-layered network exploration (Figure 2) using network propagation algorithms in order to provide new biological insights.

**Figure 2:**
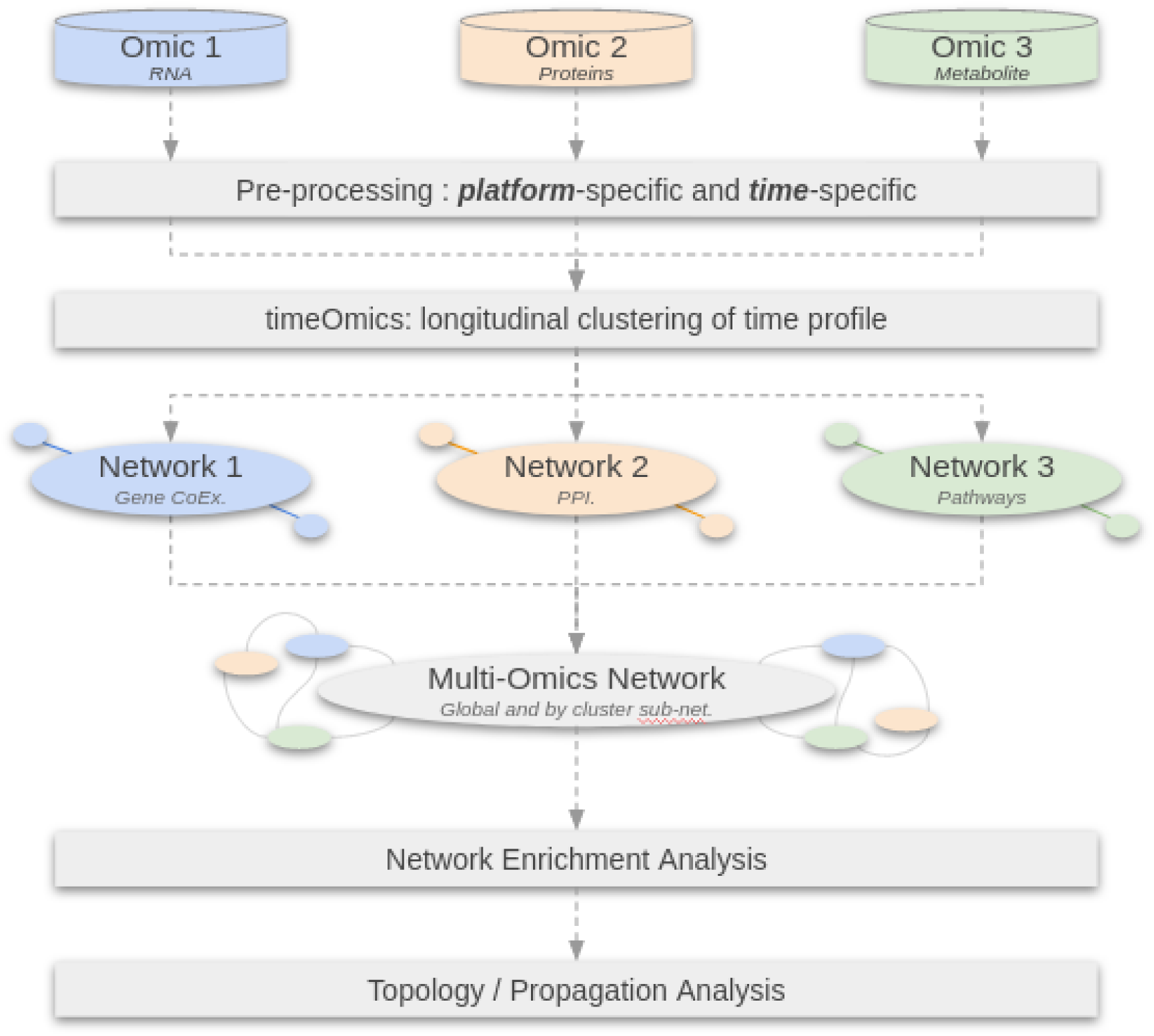
Workflow diagram illustrating the main steps of network based integration using longitudinal multi-omics data.

### 2.1 Multi-Omics longitudinal design

We define longitudinal multi-omics designs as follow. From the same biological sample, omics data are produced (RNA, proteins, etc.) at different timepoints. Raw data are processed to get (*NxP*) abundance tables by omics data type with samples in rows and molecules or biological features (RNA, proteins, metabolites, etc.) in columns. We these tables *blocks*. In this framework, there is no need to have matching timepoints between blocks because we use a modelling step to interpolate missing timepoints and even out uneven designs.

### 2.2 Pre-Processing of longitudinal multi-omics data

We assume each omics data is a raw count table resulting from bioinformatics quantification pipelines [4, 47]. Low counts are filtered and data are normalized according to the type of data in each table. We also applied a filter on time profiles and kept only molecules with the highest expression fold change between the lowest and highest point over the entire time course, as described in [3]. For each case study, we adapted these filters to take into account platform-specific dynamic range of values.

### 2.3 timeOmics: modelling and clustering of longitudinal multi-omics data

The timeOmics approach [3] was used to cluster multi-omics molecules with similar expression profiles over time. The framework is based on 2 main steps:

1. The first step uses a Linear Mixed Model Spline framework [40] to model every molecule over the time-course by taking into account the inter-individual variation. This framework tests different models and assigns the best model to each molecule according to a goodness of fit test. One of its benefits is to allow interpolation of missing timepoints and thus accommodate non-regular experimental designs with missing data.
2. The second step clusters the modelled expression profiles in groups of similar expressions over time. This is performed using various multivariate projection-based methods implemented in mixOmics [36]. With a 3-blocks omic design, we used multi-block Projection on Latent Structures (block PLS) to cluster time profiles from multi-omics datasets. Optimal number of clusters is determined by maximising the average silhouette coefficient.

The objective of these preliminary steps is to summarise each molecule with an expression pattern. This tag will then be used in the multi-omics network reconstruction to build cluster-specific sub-networks.

### 2.4 Network reconstruction

In order to build a multi-omics network, we started by building a map for each layer (genes, proteins, metabolites, etc) using a combination of both data-driven and knowledge-driven building methods. The method used is specific to the type of data and are described below. As mentioned in section 2.3, we kept the clustering information by building several sub-networks per kinetic cluster. We also built an entire network without the cluster labels.

#### 2.4.1 Data-driven network reconstruction

For gene expression data, we relied on Gene Regulatory Network inference. This class of method tries to reverse engineer complex regulatory mechanisms in the organisms and infer relationship between genes. ARACNe [27] is a co-expression-based inference algorithm which identify most likely TF - targeted genes interaction by estimating mutual information, a similarity distance between pairs of transcript expression profile. We used ARACNe algorithm on gene expression profile to infer potential TF-targeted genes interaction from the gene expression dataset.

#### 2.4.2 Knowledge-driven network

Some kind of interactions cannot be revealed by inference methods and has to be experimentally determined (for example, Protein-Protein interactions, ChIP). We then relied on reviewed interactions found in specialized databases to connect molecules of the same type and also get cross-layered interactions (binding, enzyme, regulation) From the measured molecules in the dataset, we collected all possible interactions through targeted databases. To maximize cross-layered connectivity, we also included non-measured proteins or metabolites which were directly connected to measured molecules.

##### Protein Interactions

Physical or functional Protein-Protein Interaction (PPI) is one kind of interaction that is difficult to predict and PPI network inference algorithms for MS data are still in their infancy [51]. For human proteomic data, we relied on the BioGRID database [5] which records more than 1.8 million proteins and genetics interactions from major model organisms. This database collects experimentally determined physical protein-protein interactions and also connects transcriptomics and proteomics layers with regulatory relationship (TF - gene interactions). For other model organisations, we can rely on more specialised or custom databases [21].

##### Metabolite Interactions

We used the KEGG Pathway database [19] which records a collection of manually drawn metabolic pathways representing molecular interaction, reaction and relation networks for human and other model organisms. We used KEGG to link metabolite compounds involved in the same reactions. We also connect metabolites to genes and/or proteins if they are involved in the same biochemical enzymatic reaction thanks to KEGG Orthology database that links genes to high-level functions.

The objective of this building step is to provide an entire multi-omics network composed of 3 main layers and several sub-networks specific to kinetic clusters. The next steps will focus on the analysis of these multi-omics networks.

### 2.5 Enrichment Analysis

Over representation analysis (ORA) helps to find enriched and meaningful biological insights from interacting biomolecules. This task was achieved using gProfiler2 [34], first on each kinetic cluster and then on all molecules from the entire network. We focused on the three Gene Ontology (GO) terms: Biological Process (BP), Molecular Function (MF) and Cellular Component (CC). P-values were corrected with gProfiler2 custom multiple testing correction algorithm (g:SCS) [35] and only significant terms were considered (g:SCS *<* 0.05). Size and significance of p-values distributions were compared between both clusters and entire network approaches. We also used Fisher’s combined probability test [37] for multi-omics p-values comparison.

### 2.6 Random Walk

As described in [44], in an undirected graph *G* = (*V, E*), the random walk (RW) starts from a node (*v*_0_), called *seed*, and simulate a particle that randomly moves from one node *v*_*t*_ to another *v*_*t*+1_ following the probability distribution: ∀*x, y* ∈ *V*, ∀*t* ∈ ℕ

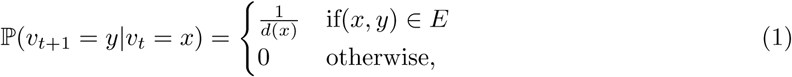

where *d*(*x*) is the degree of the node *x* in the graph *G*. Valdeolivas et al. also added the possibility to restart at the initial node to avoid dead ends in multi-layered networks. When a steady state is reached, the algorithm gives a probability score to each node of the network which represents the proximity of that node and the seed.

We then used the R package `RandomWalkRestartMH` [43] to apply random walk with restart algorithm on multi-omics network with three main purposes to guide interpretation. (1) RW can be used to identify multi-omics nodes and their interactions linked to mechanisms of interest (e.g. GO:BP). Therefore, a GO term node can be turned into a seed and RW can be performed from that starting point. Then, a sub-network with the top 25 closest nodes to that seed can be built. Naively, all significant GO term nodes were iteratively turned into seeds. We then screened sub-networks containing different types of molecules to highlight the multi-omics aspects of the integration. We applied this analysis on both kinetic cluster sub-networks and entire network. (2) RW can be used for nodes function prediction. Similar as above, unlabelled nodes can be turned into seeds. We relied on gProfiler2 annotations to identify nodes without any known functions. For an unlabelled seed, a list of ranked nodes was produced and the closest GO term node was assigned to that seed. We repeated this for the 3 GO ontologies (BP, MF, CC) on the entire network. (3) Combined to kinetic clusters, RW can locate regulatory mechanisms and find interacting nodes with different expression profiles in the entire network. Once again, each node can be turned into seed and sub-networks were built using the 10 closest nodes. Then we screened sub-networks with different cluster labels from the seeds that might reveal underlying regulatory mechanisms.

### 2.7 Data

In the following section, two published multi-omics case studies are presented. These applications have longitudinal multi-omics designs, but each block was analysed separately. We modelled these dataset with multi-layer networks to highlight the multi-omics interactions. Specific analysis steps for each example are described, along with specific databases used for each case study.

#### 2.7.1 Case Study 1: HeLa Cell Cycling study

Understanding the complex relationship between gene expression, translation product and protein abundance is the key to decrypt biological mechanisms. Genes undergo several steps of regulation before turning into proteins including transcription, translation, folding, post-translational modification and eventually degradation. [1] studied the poor correlation between mRNA and related protein levels during cell cycling regulation using triplicate measurement of mRNA expression (microarray), translation product (PUNCH-P) and protein quantification (MS) from synchronized HeLa S3 cells. Authors sampled cells during phases G1, S and G2 (Figure 3).

**Figure 3:**
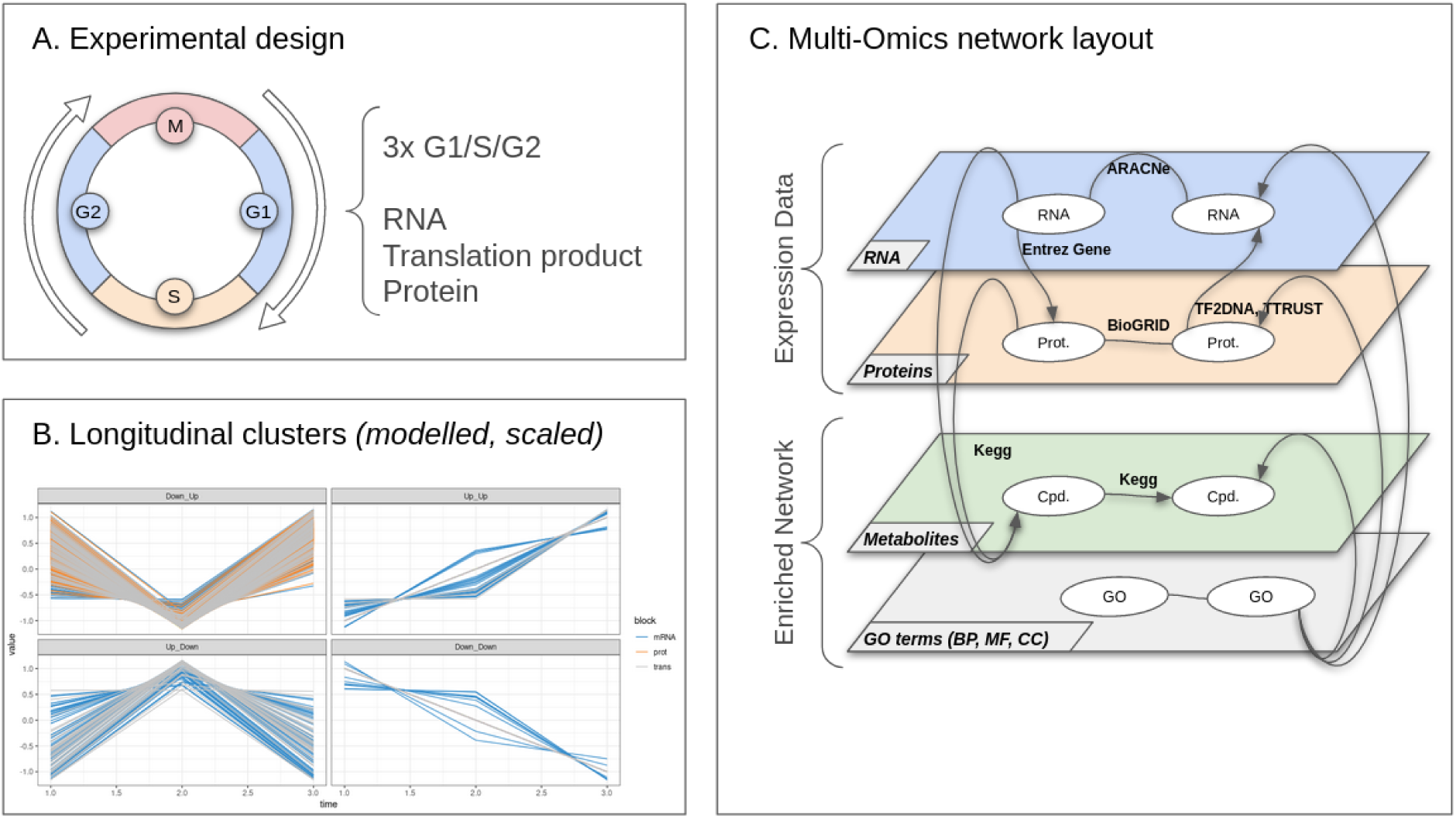
HeLa cell cycling study: overview of the analysis: (A) Experimental design: 3 samples are collected for each steps G1, S and G2. For each sample, RNA, translation products and proteins are quantified. (B) Multi-Omics longitudinal clustering: Clustered expression profiles of mRNA, translation products, and proteins. Each line represents the modelled abundance of a molecule during the cycle. 4 clusters were obtained using the timeOmics clustering approach. Cluster compositions are detailed in Table 2. (C) Multi-omics network layout: represents the connection between entities and their different types of interactions.

The authors were able to find clusters of patterns in the expression of genes and their products related to key functions of the cell cycle. These functions were up- or down-regulated at different stages. For example, *Cell division, Cytokinesis, Spindle, Chromosome segregation* and *Microtubule based movement* biological processes shared similar patterns and were down-regulated in G1/S and up-regulated in S/G2 transitions. Interestingly, multi-omics showed that a gene and its derivatives can have different expression patterns during a given time course, which highlights underlying molecular mechanisms, such as mRNA or protein degradation.

In this first case study, we intended to highlight the multi-omics interactions involved in the control of HeLa cell cycle during G1, S and G2 phases.

To do so, we filtered the RMA-normalized mRNA with a 2-Fold-Change filter. iBAQ normalized translatome products and proteins were filtered with a different 3-Fold-Change threshold because of the differences in platform-specific dynamic ranges.

We modelled the expression of every molecule with LMMS including the variations of the three replicates for each of the three timepoints. We built multi-omics clusters of expression profiles based on the direction of variation between each step. mRNA and proteins networks were built with ARACNe and BioGrid interaction databases, respectively. Protein coding and TF regulated information were used to connect these layers where UniProtID were converted into Gene Symbols (https://www.uniprot.org/uploadlists/) to ensure proper matching. We also included metabolite reactions from KEGG connected to protein enzymes to metabolites. We performed ORA with mRNA, translatome products and proteins against GO:BP, GO:MF and GO:CC terms for both clusters and entire set of molecules. Finally, we performed RW on this multi-omics network.

#### 2.7.2 Case study 2: Dynamic Maize Responses to Aphid Feeding

Maize *(Zea mays)* is one of the most productive cereal crops in the world. However, the plant is subject to numerous biotic attacks caused by herbivorous insects and it is therefore critical to understand the maize defense mechanisms in order to improve its productivity. Aviner et al. studied the dynamic of maize response to aphid feeding and they found that mutants in benzoxazinoid biosynthesis and terpene synthases genes do affect aphid proliferation.

To measure gene expression changes over time, authors exposed five two-weeks maize plants (B73) to corn leaf aphid *(Rhopalosiphum maidis)* during four days (Figure 7). They also include five control maize plants with the same growing conditions minus the exposure to aphids. During this time course, they sampled the five exposed and five controls plants at six timepoints (i.e. 2h, 4h, 8h, 24h, 48h, 96h) and conducted gene expression profiling with RNA-seq, LC-TOF-MS nontargeted metabolite quantification as well as amino-acids, phospholipids and terpenes targeted metabolite quantification.

**Figure 4:**
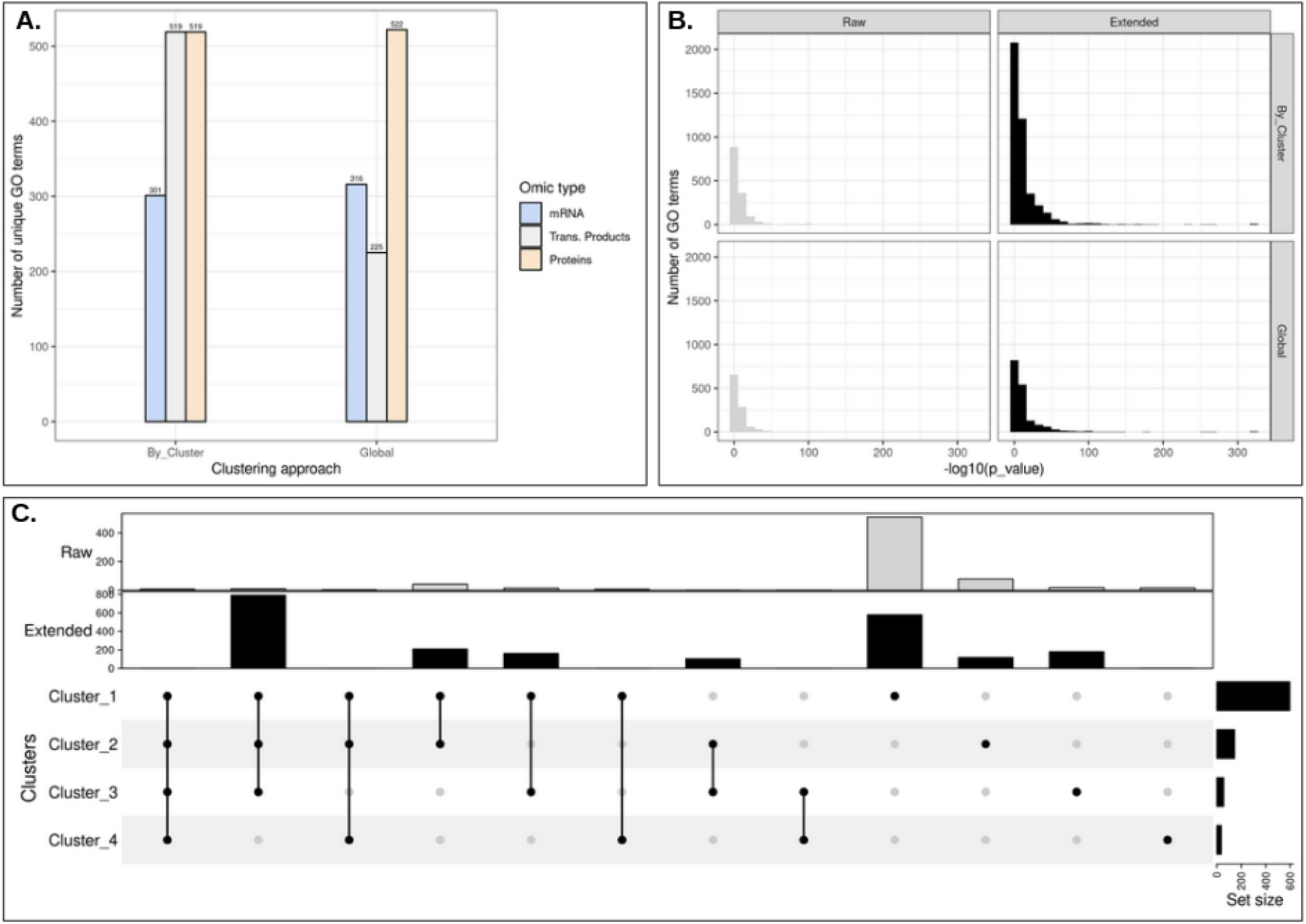
HeLa cell cycling study: Over Representation Analysis of mRNA, transcription product and protein term list. (A.) Number of distinct go terms with or without clustering approach. (B.) Histogram of p-values resulting from GO term enrichment by clustering approach. We separated the terms obtained from measured molecules (raw), and terms from with BioGRID first degree neighbours (extended). (C.) Intersection of go terms by clusters. None of the approaches seemed to offer the most enriched go terms. Nevertheless, **C**. shows that there were many terms not shared between clusters.

**Figure 5:**
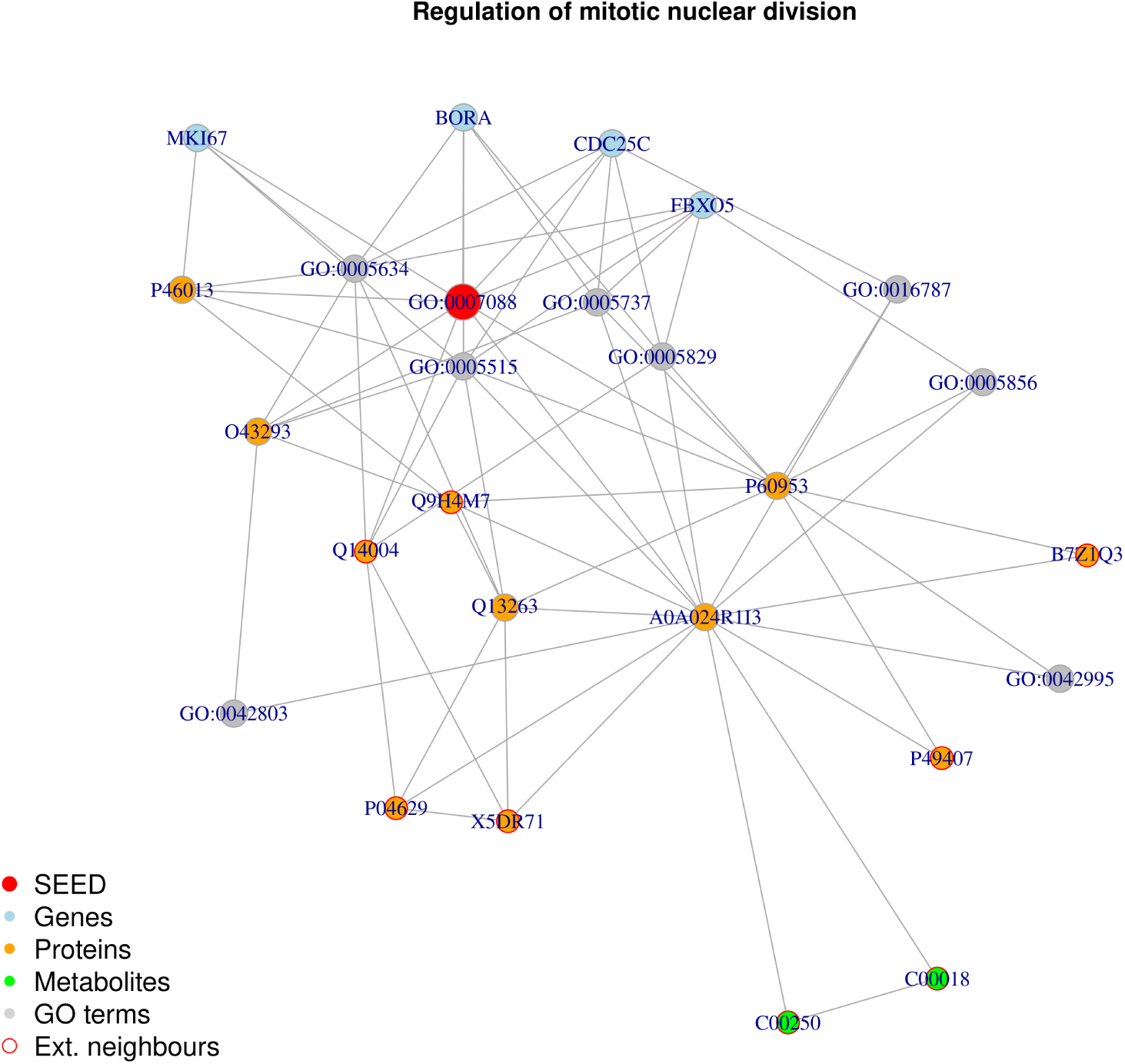
HeLa cell cycling study: Random walk with restart result from the seed GO:0007088. The top 25 results are displayed. From the seed, the algorithm is able to associate genes, proteins and go terms. Thanks to the addition of knowledge-based layers, it can also access other proteins and metabolites which were not produced in the original study (Ext. Neighbours).

**Figure 6:**
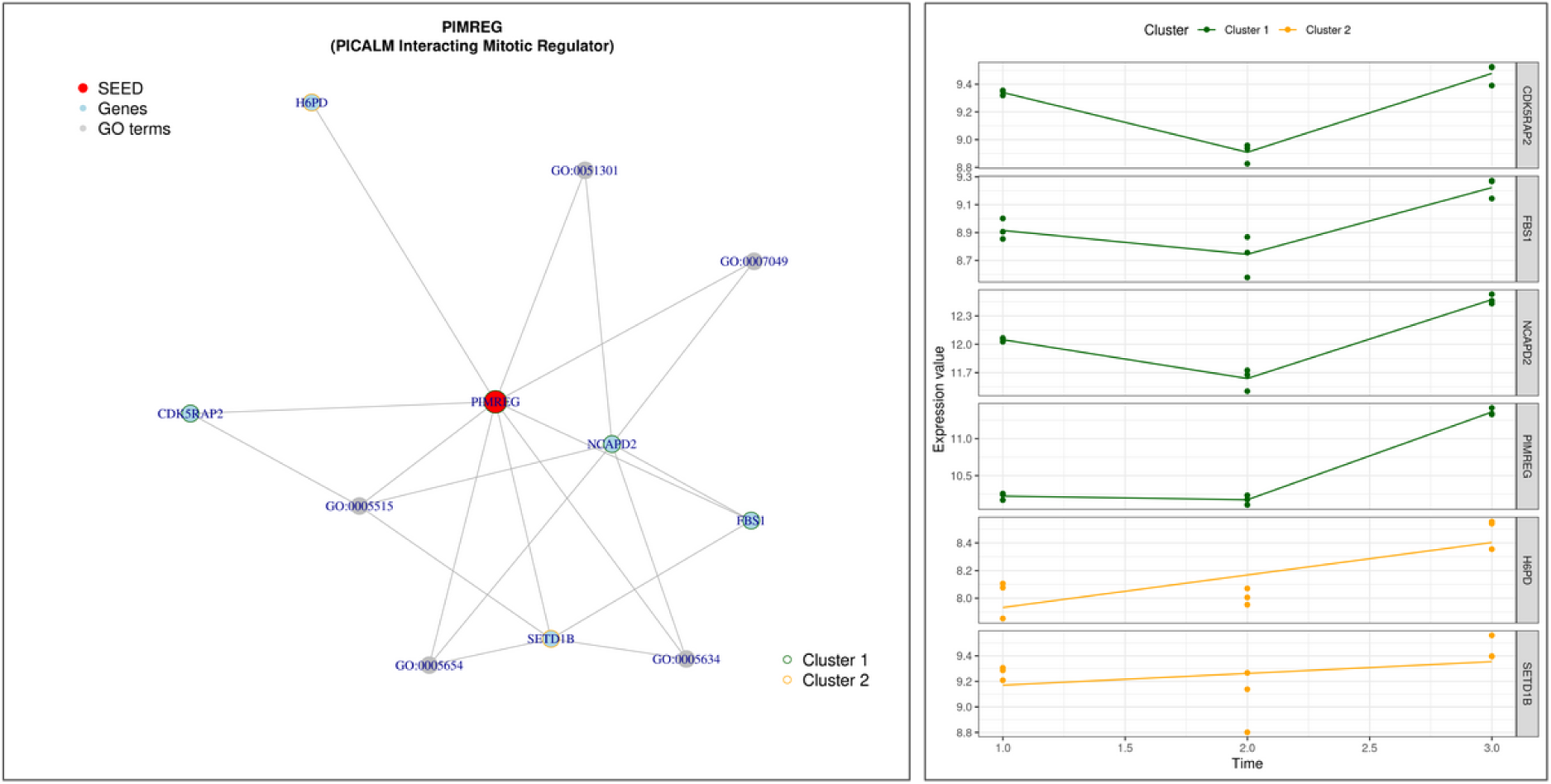
HeLa cell cycling study: Random walk with restart result analysis from the node *PIMREG*. **(A**.) Sub-network generated with top 10 RW closest nodes, colored by kinetic clusters. **(B**.) Expression profiles of measured genes from **A**.. Each line represent the modelled expression of a measured gene.

**Figure 7:**
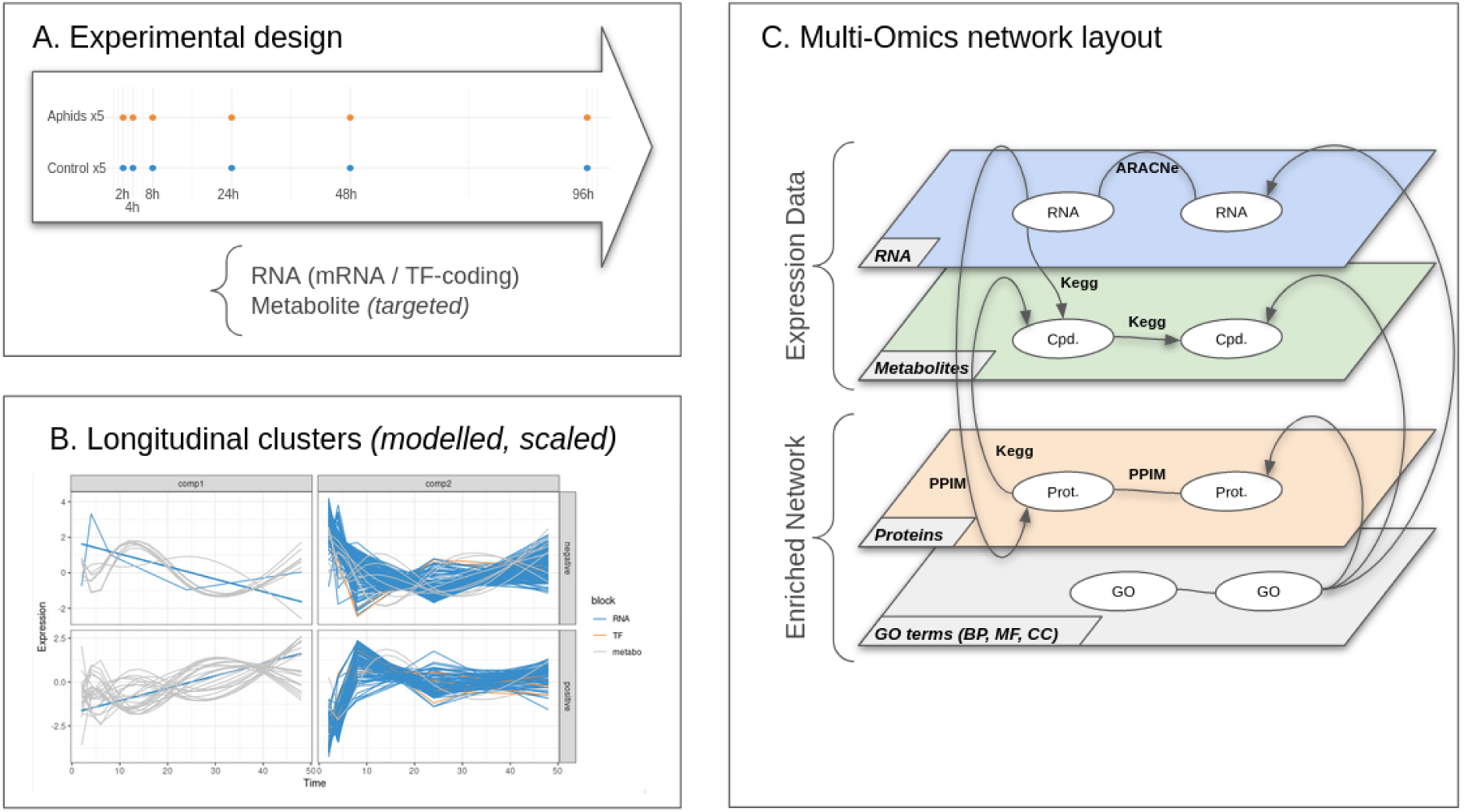
Dynamic response to maize aphid feeding study: (A) Experimental design: 5 samples are collected for each condition and for each timepoint (2h,4h,8h,24h,48h,96h). For each sample, RNA, and targeted metabolites are quantified. (B) Multi-Omics longitudinal clustering: Clustered expression profiles of mRNA, TF-coding genes and metabolites. Each line represents the modelled abundance of a molecule during the time-course. (C) Multi-omics network layout: represents the connection between entities and their different types of interactions.

In the paper, the authors focused on the genes and metabolites involved in the maize response to aphids. We intended to go a little further by addressing the complex regulatory relationships that may exist between these multi-omic actors over time.

For this example, we focused on the first five timepoints (i.e. 2h, 4h, 8h, 24h, 48h). We discarded genes which were not differentially expressed between exposed and control groups and those having an expression difference less than 2-fold over the entire time course. We split TF-coding genes from other transcripts to get a 3-omics-like experimental design. Finally, we discarded nontargeted metabolite as they were not annotated and performed longitudinal clustering with multiple block PLS. For network reconstruction, we used ARACNe on genes and TF. We used the Protein-Protein Interaction database for Maize (PPIM) [53] to identify protein-coding genes and extracted direct neighbor interactions. PPIM contains more than 2.7 millions interactions between protein, TF and gene interactions. These interactions are either predicted or experimentally determined from public databases such as UniProt [7], BioGrid [5], DIP [46], IntAct [31] and MINT [24] or using to text mining. We mainly focused on validated interactions and predicted ones qualified as high-confidence interactions that represents 155,845 interactions with top 5% highest decision scores [53]. Then we connected the measured metabolites and those involved in the same reaction to the genes/proteins using KEGG. We performed ORA with genes, TF and proteins on GO:BP, GO:MF and GO:CC terms. Finally, we performed RW on this multi-omics network.

## 3 Results

### 3.1 Case study 1: HeLa Cell Cycling study

In this example, we studied the HeLa cell cycling from Aviner et al. 1. This dataset was composed of 3 omic layers and 3 timepoints.

The network was composed of 4 layers: a gene layer build from mRNA expression, a PPI layer build from measured proteins and BioGRID known interactions, a metabolite layer from KEGG pathways and GO term layer from enrichment analysis.

#### 3.1.1 Pre-processing, modelling and clustering of time profile

HeLa cell dataset was assembled into a single table focused on proteins. After missing value removal, this dataset was composed of 6,785 mRNA, 4,102 translation products and 5,023 proteins (Table 1). 448 mRNA, 2,672 translation products and 4,295 proteins remained after the Fold-Change filtering step. Finally, once all molecules were modelled over time and *noisy* profiles were removed, 446 mRNA, 2,318 translation products and 4,237 proteins remained.

**Table 1:** HeLa cell cycling study: Initial number of mRNA, translation products and proteins, and remaining molecules after Fold-Change and noisy modelled profile filtering. A threshold of 2-FC was applied to mRNA and 3-FC to both proteins and translation products.

|  | Raw Counts | Fold Change | LMMS filter |
| --- | --- | --- | --- |
| mRNA | 6,785 | 448 | 446 |
| Trans. Products | 4,102 | 2,672 | 2,318 |
| Proteins | 5,023 | 4,295 | 4,237 |

#### 3.1.2 Time profile clustering

Once all the molecules were modelled over time and noisy profiles were removed, remaining expression profiles were clustered according to their differential expression value between two timepoints. This clustering resulted in 4 clusters (Table 2) with a silhouette coefficient of *s*_*sil*_ = 0.75. We compared this clustering with the *timeOmics* approach with 4 clusters for similar parameters. *timeOmics* resulted in a lower silhouette coefficient (*s*_*sil*_ = 0.63). In the first approach, Cluster 1 included the largest number of molecules (*n* = 4, 443). It is characterized by a decrease between the G1 and S phases and then an increase between S and G2/M. This seemed to be the main kinetic pattern for proteins since it contained the majority of them (95%). Cluster 2 (*n* = 1, 444), included the largest amount of translation products (54%) and it was characterized by an increase in expression from the first to the last step. Cluster 3 (*n* = 958) showed an opposite pattern compared to the cluster 2 with a decrease across the overall time course. Finally Cluster 4 (*n* = 156) included the least number of molecules and no protein appeared to follow an increase and decrease pattern.

**Table 2:** HeLa cell cycling study: Clusters composition

|  | mRNA | Translation products | Proteins |
| --- | --- | --- | --- |
| Cluster 1 | 233 | 187 | 4,023 |
| Cluster 2 | 81 | 1,244 | 119 |
| Cluster 3 | 35 | 828 | 95 |
| Cluster 4 | 97 | 59 | 0 |

#### 3.1.3 Multi-layered network reconstruction

The first layer to be reconstructed was the gene inference network from the mRNA. We used the ARACNe algorithm to build a network by kinetic cluster but also to build a entire network composed of all the mRNA. Suppl. Table 1 shows statistics about the sub-networks such as the number of connected / disconnected nodes and edges.

The second layer was the PPI network. Proteins were connected to each other using the BioGRID interaction database. As for the genes, PPI sub-networks were built for each kinetic cluster but we also built a entire PPI network composed of all the proteins. In addition, we included BioGRID proteins which were directly connected to the measured ones (first degree neighbours). Number of nodes and edges are detailed in Suppl. Table 1.

These first 2 layers were combined thanks to 2 types of links. First, protein-coding information linked 126 genes to their corresponding proteins. Second, TF-regulated information from TF2DNA and TTRUST databases linked 57 proteins to 403 genes (16,846 interactions). In addition, KEGG pathway database was also used to link protein enzymes to metabolite reactions. This new layer was composed of 1,595 metabolites connected to 2,213 proteins (12,694 interactions).

#### 3.1.4 Enrichment Analysis

Over representation analysis (ORA) was performed on mRNA, translation products, proteins and first degree connected proteins against GO:BP, GO:MF and GO:CC. This approach was achieved both for each kinetic cluster and the entire network. The most significant GO terms found in clusters were strongly related to cell development *RNA processing (GO:0006396), ribonucleoprotein complex biogenesis (GO:0022613), cellular component organization or biogenesis (GO:0071840)* or *RNA splicing, (GO:0008380)*. Full results of the ORA are detailed in Suppl. Table 2. With the largest number of molecules, proteins gathered the most significant GO terms. Table 3 details the number of enriched GO terms by omic types. Enriched terms were splited by ontologies. mRNA collected the fewest enriched terms in the 3 ontologies, followed by translation products and proteins. The first degree neighbours (Extended proteins) collected the most terms, as they have the highest number of nodes in the network. In figure 4, we compared the number of unique GO terms and the distribution of P-value between the global or cluster approach. With the exception of translation products which had more enriched term in clusters than the entire set, both approaches seemed to have a similar number of unique GO terms but some terms *DNA replication (GO:0006260), mitotic cell cycle (GO:0000278)* were shared between clusters. In addition the clustering approach gave much more significant terms than the global approach with the extended proteins. There was no significant difference in the p-value distributions however the clustering approach results in smaller p-values. In the Fig. 4.**C** some GO terms were exclusively found in certain clusters such as: *regulation of cell projection organization (GO:0031344)* and *nervous system development (GO:0007399)* in Cluster 1, *spindle midzone assembly (GO:0051255)* and *meiotic chromosome separation (GO:0051307)* in Cluster 2, *mitotic G2/M transition checkpoint (GO:0044818)* and *regulation of DNA endoreduplication (BP) GO:0032875* in Cluster 3, *negative regulation of gene expression, epigenetic (GO:0045814)* and *regulation of chromatin silencing (GO:0031935)* in Cluster 4. Moreover some terms were also only found in the entire set such as *chromosome condensation (GO:0030261), regulation of cytokinesis (GO:0032465)* and *histone H3 acetylation (GO:0043966)*.

**Table 3:** HeLa cell cycling study: Number of unique significant enriched GO terms by omics type and by ontology.

|  | GO:BP | GO:CC | GO:MF |
| --- | --- | --- | --- |
| mRNA | 255 | 82 | 54 |
| translation products | 331 | 124 | 70 |
| Proteins | 353 | 128 | 69 |
| Extended Proteins | 1490 | 390 | 280 |

Finally, significant GO terms were added in the multi-layer network as another new layer and were connected to genes, proteins and metabolites. This last building step resulted in a multi-omics network composed of 4 layers. Table 4 describe network composition with the different type of node for each layer.

**Table 4:** HeLa cell cycling study: Multi-Omics network composition. Number of nodes by layer.

|  | Gene | Proteins | Extend Prot. | Metabolite | GO terms |
| --- | --- | --- | --- | --- | --- |
| Cluster 1 | 233 | 1813 | 22387 | 1585 | 1576 |
| Cluster 2 | 81 | 53 | 4726 | 591 | 1325 |
| Cluster 3 | 35 | 50 | 4205 | 559 | 1328 |
| Cluster 4 | 97 |  |  |  | 441 |
| All | 252 | 1916 | 22582 | 1595 | 1590 |

#### 3.1.5 Random Walk

##### Molecules involved in specific mechanisms

The first purpose of RW applied to multi-omics network was to identify molecules involved in a specific mechanisms. Naively, each of the 2,279 significantly enriched GO terms, (union of GO terms found in cluster and the entire network), can be a mechanism or a function of interest. Each of these GO terms was then turned into a seed. RW from each seed were used to generate sub-networks the 25 top closest nodes. In order to focus on multi-omics integration, we only focused on seeds that reached both genes and proteins. In a second step, we also included seeds that reached both genes, proteins and metabolites. Table 5 describe the number of GO terms (BP, MF, CC) where these conditions were met. 16 unique terms were linked to metabolite nodes (12 BP, 4 MF). Some terms were shared between clusters. These terms encompassed functions related to cell development such as *histone lysine methylation GO:0034968* or *regulation of mitotic nuclear division (GO:0007088)*. In Figure 5, we illustrate in detail the multi-omics interactions leading to the activation of this last biological process *(GO:0007088)*. More general terms were also found: *positive regulation of cellular metabolic process (GO:0031325), mRNA catabolic process (GO:0006402)*. Some MF terms seemed also related to cell development *anaphase-promoting complex binding (GO:0010997), S-adenosylmethionine-dependent methyltransferase activity (GO:0008757)*. We also found such terms as *response to ionizing radiation (GO:0010212)* which could be an artifact of mass-spectrometry analysis. On the other hand, besides the term *histone methyltransferase activity (H3-K27 specific) (GO:0046976)* other shared terms seemed to be rather generic : *methyltransferase activity (GO:0008168)* or *regulation of signal transduction (GO:0009966)*.

**Table 5:** HeLa cell cycling study: (A.) Number of seeds that can reach both genes and proteins. (B.) Same as A but with the addition of metabolites.

| **A.** | GO:BP | GO:CC | GO:MF | **B.** | GO:BP | GO:MF |
| --- | --- | --- | --- | --- | --- | --- |
| Cluster 1 | 127 | 57 | 46 | Cluster 1 | 7 | 1 |
| Cluster 2 | 98 | 59 | 35 | Cluster 2 | 2 |  |
| Cluster 3 | 57 | 23 | 25 | Cluster 3 | 1 | 2 |
| All | 187 | 82 | 56 | All | 9 | 2 |

##### Function Prediction

The second purpose of RW was to predict the function of unannotated nodes (genes, proteins). In this perspective, 6 genes and 24 proteins were unannotated according to gProfiler. Each of these nodes was then turned into a seed. We then assigned to each node the closest GO term. Results of function prediction annotation are detailed in the Suppl. Table 3. The first biological processes which emerged for almost every unannotated nodes were *cell cycle (GO:0007049)* and *cell division (GO:0051301)* with *protein binding (GO:0005515)* or *RNA binding (GO:0003723)* molecular functions and were related to the nucleus *(GO:0005634)*. Another interesting example was the uncharacterized protein *DKFZp781F05101*, which should play a role in the nucleosome formation *(GO:0006334)* and in particular in this study those containing the histone H3 variant CENP-A to form centromeric chromatin *(GO:0034080)*.

##### Intersection between cluster

The last objective of the RW was to be combined with the kinetics cluster in order to identify connected molecules with different expression profiles. All the nodes representing a measured molecule were transformed into a seed. From the 2,362 seeds, 438 were connected to molecules from other clusters We kept 47 seeds that can also reach GO term nodes. These results are detailed in Suppl. Table 3. We identified many biological processes terms related to the cell development. For example, the node labelled *PIMREG* was a protein coding gene belonging to Cluster 1. With RW, it was linked to Cell Division (GO:0051301), Cell Cycle (GO:0007049) and other GO:BP terms. PIMREG is a mitotic regulator which takes part in the control of metaphase-to-anaphase transition. It was linked to *SETD1B* (Cluster 2), an histone methyltransferase that tri-methylates histone H3 at Lysine 4 and contributes to epigenetic control of chromatin structure and gene expression. This is an active marker of up-regulated transcription and this may explain why the expression of the PIMREG gene is increasing between phases S and G2/M. In Figure 6 *PIMREG* is also directly connected to the *CDK5RAP2* gene, which has the same expression profile as *PIMREG* and plays a role in the formation and stability of microtubules, essential to the cell development.

### 3.2 Case study 2: Dynamic Maize Responses to Aphid Feeding

#### 3.2.1 Pre-processing, modelling and clustering of time profile

The maize dataset included a gene expression assay composed of 41,716 transcripts identified using the B73 maize reference genome (AGPv3.20), with 2,254 transcripts identified as TF-coding genes and a targeted metabolite assay with 45 molecules measured. 1,606 maize genes, 123 TF and all metabolites were differentially expressed between aphid and control groups for at least 1 time points. 331 genes, 36 TF and 29 metabolites were filtered after the 2 Fold-Change filtering step. Once all the molecules were modelled and *noisy* profiles were removed 1,208 genes, 31 TF and 29 metabolites remained in the exposed group (Table 6).

**Table 6:** Dynamic response to maize aphid feeding study: Initial number of genes, TF-coding genes and metabolites and after each pre-processing step *(DE: Differentially Expressed molecules; FC: molecules above 2 Fold-Changes, LMMS filter: number of molecules after linear model filtering)*.

|  | Raw count | DE filter | FC filter | LMMS filter |
| --- | --- | --- | --- | --- |
| Genes | 39,462 | 1,606 | 1,295 | 1,208 |
| TF-coding genes | 2,254 | 123 | 75 | 42 |
| Metabolites | 45 | — | — | — |

#### 3.2.2 Time profile clustering

Next step was the clustering with timeOmics using genes, TF-coding genes and targeted metabolites. This resulted in optimally 4 clusters (*s*_*sil*_ = 0.82). Table 7 gives the cluster composition. Clusters 1 and 2 recorded most of the molecules (42%; 40%). These were mainly composed of linear models representing an increase (cluster 1) or a decrease (cluster 2) in the time course. Cluster 3 (7%) was characterised by a rapid decrease followed by a slight increase from *t* = 24*h*. Cluster 4 (11%) showed the opposite of cluster 3 with a rapid increase followed by a relatively steady state.

**Table 7:** Dynamic response to maize aphid feeding study: number of gene, TF-coding genes and targeted metabolites per cluster identified with multi-block PLS longitudinal clustering

|  | Cluster 1 | Cluster 2 | Cluster 3 | Cluster 4 |
| --- | --- | --- | --- | --- |
| Genes | 496 | 495 | 84 | 133 |
| TF-coding genes | 18 | 15 | 6 | 3 |
| Metabolites | 24 | 7 | 4 | 10 |

#### 3.2.3 Multi-layered network reconstruction

The first layer to be reconstructed was the gene inference network from the mRNA and TF using ARACNe. We used unmodelled filtered data from aphid and control samples to build this first layer. As the first case study, we build sub-network for each cinetic cluster and a global network composed of all the mRNA/TF. Networks composition is detailed in Suppl. Table 4.

The second layer was the PPI network. Filtered PPIM database was used to build protein-protein interaction network from measured protein-coding genes. We also included proteins which were directly connected to the measured protein coding genes (first degree neighbours). Gene kinetic cluster subnetworks and global network were extended with PPI interactions.

The third network contained metabolic reactions. Metabolites converted to KEGG Compound ID were linked to each other with KEGG Pathways. Metabolites involved in the same metabolic reactions were also included in this layer. As for genes and proteins, kinetic cluster sub-networks and entire network were extended to connect genes and proteins to metabolites. Composition of multi-omics networks is detailed in Table 8.

**Table 8:** Dynamic response to maize aphid feeding study: Multi-Omics network composition. Number of nodes by layer.

|  | Cluster 1 | Cluster 2 | Cluster 3 | Cluster 4 | Entire Network |
| --- | --- | --- | --- | --- | --- |
| Genes | 514 | 510 | 90 | 136 | 1250 |
| Ext. Proteins | 1473 | 2000 | 176 | 1102 | 2545 |
| Metabolites (measured) | 17 | 12 | 14 | 10 | 38 |
| Metabolite (KEGG reactions) | 77 | 85 | 65 | 46 | 250 |
| GO terms | 121 | 131 | 57 | 100 | 148 |

**Table 9:** Dynamic response to maize aphid feeding study: Over Representation Analysis of mRNA term list by cluster. **(A**.) Number of significant GO terms per cluster and entire network with genes. **(B**.) Same as **A**. with PPIM first degree neighbours.

| **A.** | GO:BP | GO:CC | GO:MF | **B.** | GO:BP | GO:CC | GO:MF |
| --- | --- | --- | --- | --- | --- | --- | --- |
| Cluster 1 | 4 | 5 | 17 | Cluster 1 | 123 | 14 | 78 |
| Cluster 2 | 7 | 6 | 5 | Cluster 2 | 124 | 627 | 586 |
| Cluster 3 | - | 1 | - | Cluster 3 | 19 | 1 | 39 |
| Cluster 4 | 5 | - | 5 | Cluster 4 | 100 | -14 | 569 |
| Entire network | 37 | 5 | 31 | Entire Network | 122 | 530 | 104 |

#### 3.2.4 Enrichment Analysis

ORA was performed on genes and proteins against GO:BP, GO:MF and GO:CC. This approach was achieved both for each cluster and combined network. We first performed ORA on measured molecules only and with PPIM’s first-degree neighbours. All the results of ORA analysis are detailed in Suppl. Table 5. Cluster 1 with more genes gathered the largest number of significant terms. Terms specific to this cluster were molecular functions related to *nucleotide bindings (GO:0035639, GO:0032555, GO:0017076, GO:0032553)*. Other clusters grouped terms related to bindings or kinase activity *(GO:0043546, GO:0043424)*. Figure 8 represents the number of significant enriched GO terms and their p-value distribution. Measured molecules tended to produce more significant terms when ORA was performed on the entire set of genes rather than with the clustering. Nevertheless, with PPIM’s first-degree neighbours, there were more significant term in the clusters than in the entire set. Neighbours also gave smaller p-values. Lastly, ORA identified unique terms per cluster, not found in the entire set, and vice versa. Therefore, we found the terms mentioned for Cluster 1. Concerning the unique terms in the entire set, they included terms corresponding to *defence response to other organism (GO:0098542)* or other biological processes that can be linked to defence mechanisms such as *lignin metabolic process (GO:0009808), interspecies interaction between organisms (GO:0044419), cellular response to chemical stimulus (GO:0070887)* or *transmembrane transport (GO:0055085)*. Stranger, *defense response to oomycetes (GO:0002229)* was also significantly enriched in the entire set.

**Figure 8:**
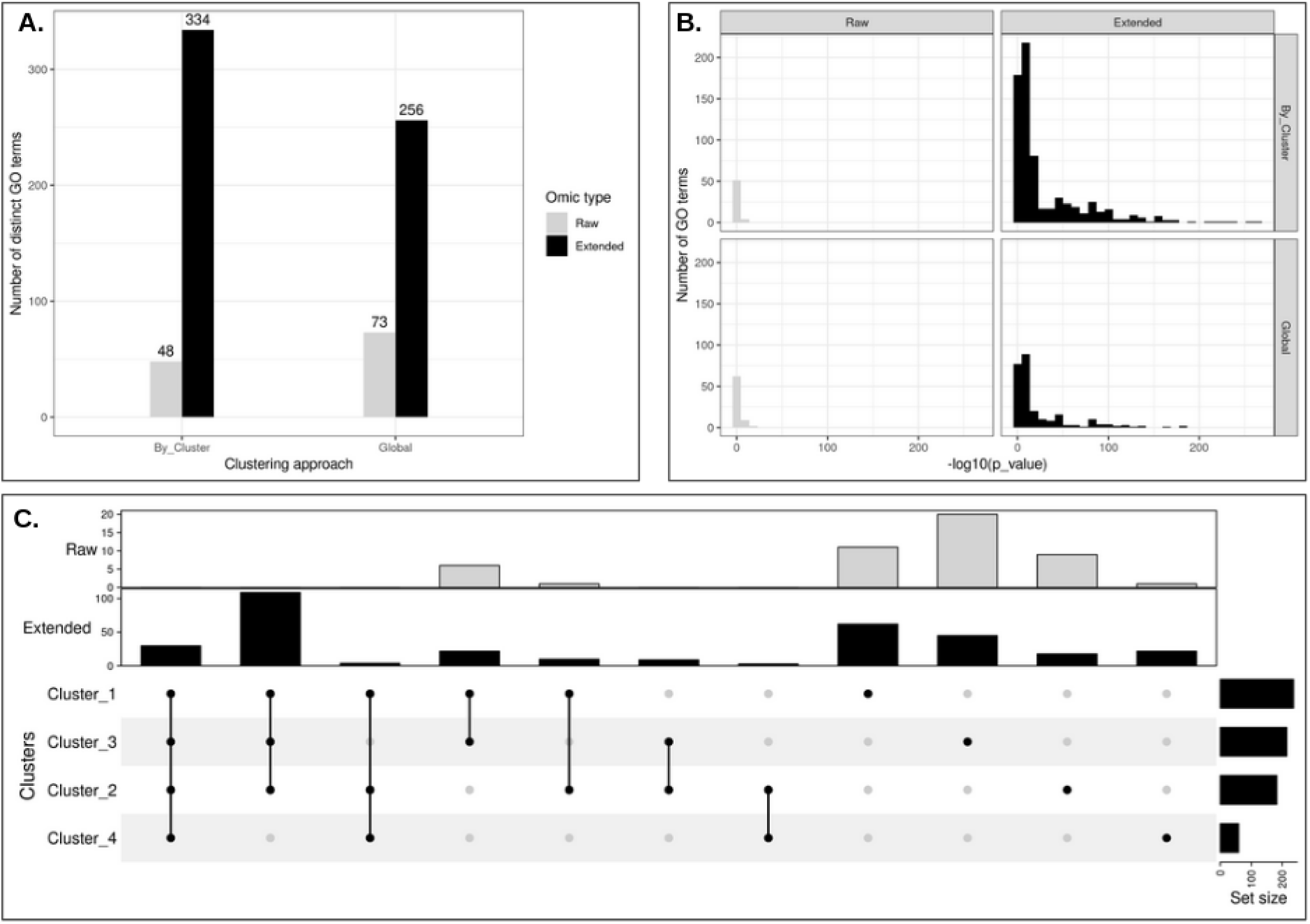
Dynamic response to maize aphid feeding study: Over Representation Analysis. (A.) Number of distinct go terms with or without clustering and by approach. (B.) Histogram of p-values resulting from GO term enrichment. (C.) Intersection of go terms by clusters

#### 3.2.5 Random Walk

Like the HeLa cell cycling study example, RW was used on the multi-omics network with 3 purposes: Identify multi-omics molecules linked to mechanisms (GO term nodes as seed), function prediction (unlabelled nodes as seeds), and identify regulatory mechanisms with nodes between kinetic clusters.

##### Molecules involved in specific mechanisms

Each of the 89 significantly enriched GO terms was iteratively turned into a seed and subnetworks with the top 25 nodes were buid. Within a multi-omics perspective, only 3 seeds were able to reach at least one gene and one metabolite: *molybdopterin co-factor binding (GO:0043546), cell surface receptor signaling pathway (GO:0007166)* and *phenypropanoid metabolic process (GO:0009698)*. The latter is specific to plants and gathers information on cell development and defence mechanisms. *Phenypropanoid metabolic process* was found in cluster 2 and 4 as well as in entire network approach. Part of the shikimate pathway that allows the production of aromatic amino acids in plants, the phenylpropanoid pathway is known to be activated under abiotic stress including salinity, drought, heavy metals, pesticides, ultraviolet radiation and temperature stress [38]. In the entire network, this node reached the metabolites *L-Tyrosine (C00082)* and *L-Phenylalanine (C00079)* which were both aromatic amino acids measured in the initial study. In addition, it reached non-measured metabolites such as *trans-Cinnamate (C00423)* and *4-Coumarate (C00811)* which were connected to *Tyrosine* and *Phenylalanine* respectively. These connections were enabled by the L-phenylalanine ammonialyase and L-tyrosine ammonia-lyase reactions in which both measured and non-measured compounds were involved. In this same sub-network, we obtained several iso-enzymes involved in these reactions such as pal1,5,6,9. Interestingly, several gene models without functional annotation are connected to the network (GRMZM2G063679, GRMZM2G087259). RW from *Phenypropanoid metabolic process* on clusters 2 and 4 networks also returned *Tyrosine* and *Tryptophan* since they were classified in clusters 2 and 4 respectively and were involved in the same metabolic reaction. It also reached genes connected to similar pathways in both clusters: *dhurrin biosynthesis* acting as a plant defense compound and also *suberin monomers, cuticular wax* and *cutin biosynthesis* for plant development. Specificaly to cluster 4, one protein (GRMZM2G164036) was linked to DIBOA-glucoside biosynthesis, a benzoxazinoids dervived from indole that is a volatile compound used in parasitic defence [11].

##### Function prediction

Unlabelled gene nodes (*n* = 96) were turned into seeds and the closest GO terms were assigned to these nodes. 347 unlabelled nodes were disconnected from the network therefore it was not possible to apply the algorithm for these ones. These results are detailed in Suppl. Table 6. Many seeds were connected to various biosynthesis process such as *cinnamic acid (GO:0009800), ceramide (GO:0046513), galactolipid (GO:0019375)*. We found unlabelled genes linked to *defense response processes (GO:0006952)* or *jasmonic acid metabolic process (GO:0009694)* which is an hormone that can be produce during biotic or abiotic stress. Other nodes were connected to various molecular functions such as *hydrolase, chitinase, methyltransferase or bindings*.

##### Cluster intersection

All measured nodes were turned into seeds and sub-neworks were build with the to 10 highest scored nodes from the entire network. 678 seeds were able to reach both nodes with different assigned kinetic cluster and GO term nodes. Results are detailed in Suppl. Table 6. We found molecular functions corresponding to different enzymatic activities (ATP binding, kinase, oxidoreductase). Less frequently, we observed a *toxin activity (GO:0090729)* for the *rip2* gene (GRMZM2G119705), also connected to *defense response (GO:0006952)*. In Figure 9, gene *AC209374*.*4 FG002*, assigned to cluster 2, were indirectly connected to nodes from both clusters 1, 3 and 4. They were linked to *DNA-binding transcription factor activity (GO:0003700)*. This regulatory mechanisms may be explained by the opposite expression profile as illustrated in Fig. 9.B.

**Figure 9:**
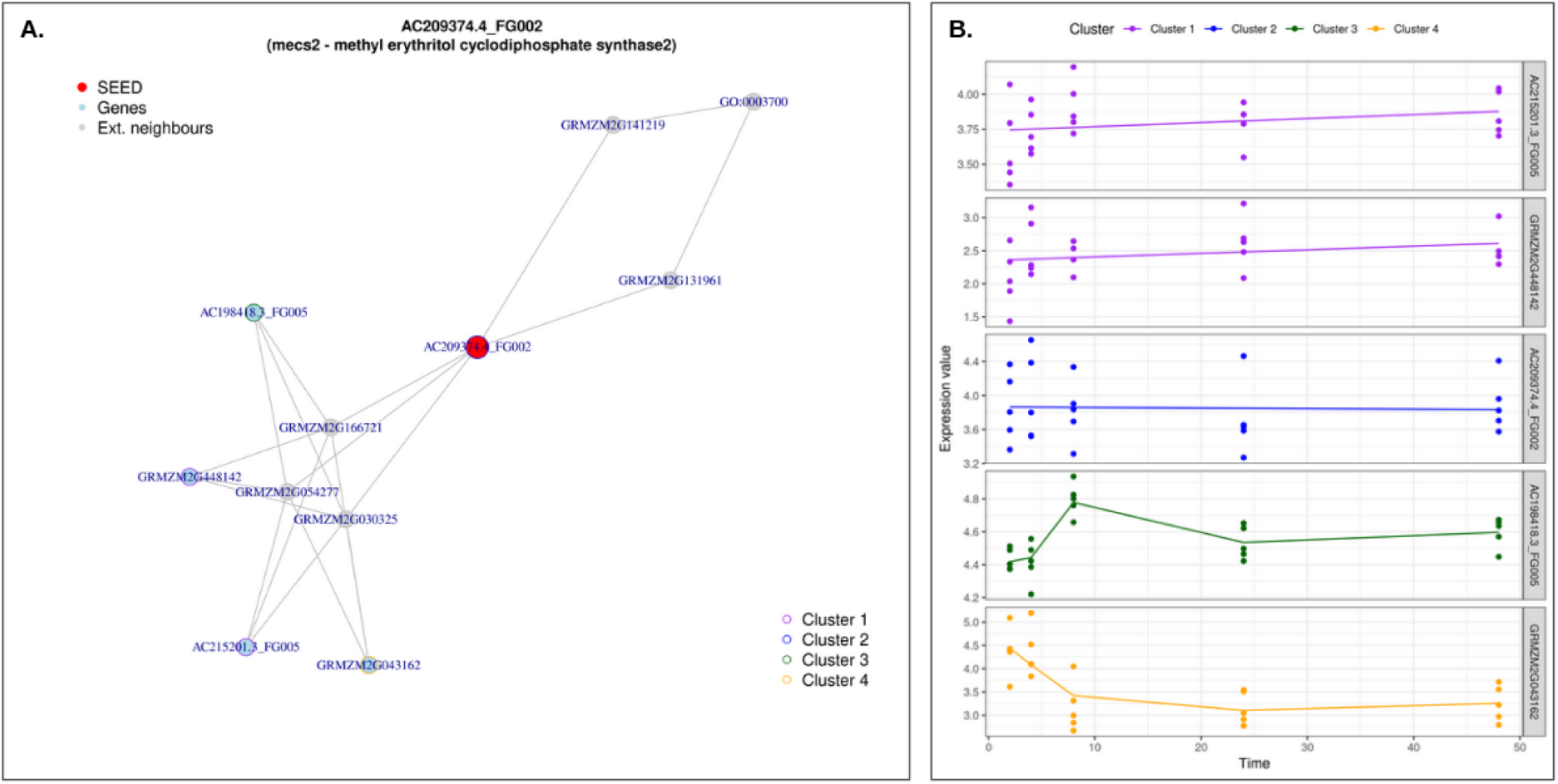
Dynamic response to maize aphid feeding study: Random walk with restart result analysis from the node *AC209374*.*4 FG002*. **(A**.) Sub-network generated with top 10 RW closest nodes, colored by kinetic clusters. **(B**.) Expression profiles of measured genes from **A**.. Each line represent the modelled expression of a measured gene.

## 4 Discussion

Improvements and cost reductions in high-throughput omics technologies have resulted in the emergence of increasingly complex experimental designs that combine both multi-omics data and longitudinal sampling from the same biological samples. These complex designs are implemented to provide a comprehensive view of biological systems in order to identify complex relationships between omics layers and answer various biological questions such as disease subtyping, biomarker discovery, models and predictions [12, 16, 29, 33]. Appropriate integration of the multiple omics profiled must be performed to fully take advantage of the interactions between omics and their interdependencies. The challenge of multi-omics integration is to offer interpretation guidelines adapted to the multiplicity and heterogeneity of designs (data types, number of conditions, samples, timepoints, etc.). In order to identify cellular mechanisms with multi-omics regulation, we propose to use a combination of both data-driven and knowledge-driven integration methods from longitudinal multi-omic data to highlight interactions not only between molecules of the same biological type but also between omic layers linked to cellular mechanisms.

### Multi-omics network integration in literature

Multi-omics networks are the key to illustrate topological relationships between different molecular species [23]. The HetioNet network is currently one of the most impressive multi-omics networks. Composed of 18 different types of nodes (genes, SNPs, proteins, compounds, tissues, diseases) and 19 different types of edges between these layers [13]. Its main objective is to provide a natural representation of the human system hierarchy and to propose genetic-disease association prediction. Although very comprehensive, some relationships evolve in time and the addition of these dynamic interactions as well as specific expression data should be very valuable to study complex diseases mechanisms. On a smaller scale, MetPriCNet [48] is a 3-level network composed of genes, metabolites, diseases and associations between each cross-layer pair. Its objective is to prioritize candidate metabolites for known diseases or phenotypes. Although very useful for therapeutic target prediction, these networks should be connected to expression data to further address the needs of personalized medicine. In another paradigm, OmicsNet [52] offers to build customized multi-omics networks from a list of molecules of interest (genes, proteins, metabolites, TF, miRNA). It does not provide association prediction but the connection between the nodes is ensured by several databases (STRING, miRBase, KEGG, ENCODE, etc.). Its main advantage is to propose in a web browser a user friendly 3D representation of the resulting network. Like other networks, it is specifically designed for the human organism and still too few integration networks target other model organisms.

### Interpretation guidelines for multi-omics networks: Advantages and Limitations

Ranging from expression modelling over time, multivariate clustering to network reconstruction, in the following section we discuss the pros and cons of the choices made at each stage of this methodology.

1. **Experimental design:** Each experimental design is different, both in type of data, number of replicates, condition and timepoints. Regardless of the normalization step specific to each technology, our previous timeOmics framework can indeed accommodate several design variations to identify highly associated temporal profiles between multiple and heterogeneous datasets. We already discussed the advantage of using PLS and block PLS over other longitudinal clustering methods and made some recommendations where timeOmics works the best (i.e. 3 blocks of data, 5-10 not too widely spaced timepoints). If there are too much timepoints or inter-individual variability is too high the paper also give guidelines.
2. **Network inference**: We relied on ARACNe algorithm for gene inference network. More mathematically advanced alternatives exist. [50] recently applied Gaussian graphical model with graphical lasso to infer gene interactions in 15 specific types of human cancer using RNA-Seq expression data from The Cancer Genome Atlas. Nevertheless, ARACNe is a well-cited, reliable coexpression-based inference method and it provides an easy to use implementation. As mentioned by [9], the inference problem is NP-complete and only the combination of the results of several methods would bring us closer to the truth. Moreover, the choice of the inference method can be directed by the type of omics data. With microbiome abundance data, co-occurrence network inference is often applied to identify synergism or competition relationships between species and their environment [26, 28].
3. **Knowledge-based interaction**: The proposed hybrid multi-omics networks were composed of expression data and knowledge links. Therefore, known links can be limited by incomplete databases. In the first case study, we mixed BioGRID, TTRUST and TF2DNA to connect proteins to genes. However, more specialised databases could also be added to include more links. In the second case study, maize is not as well-resourced as humans and PPIM covers all known to date interactions for this organism. In addition, these links represent potential interactions that may append in the studies and the networks are referred to as *maps*. An additional integration step would consist in constructing a dynamic signal flow where the directionnality and intensity of the interactions would be represented in the form of flows or pathways. Unfortunately this type of representation requires a time-series design using precise multiple doses of stimulation under the same condition [49].
4. **Multi-omics interpretation**: The use of clusters enables kinetic-specific enrichment analyses to be performed. These results can be compared with those obtained from the entire network. Both approaches tended to provide the same number of significant terms are instead complementary. It was possible to identify terms unique to each approach and terms only found in a particular cluster. Addition of non-measured molecules in ORA provided more significant terms with lower p-values. However, these extra molecules were highly shared between cluster or were linked to similar functions. Thus it did not bring cluster specific enrichment. Finally, although random walk propagation analysis is the state of the art for association prediction, its application on heterogeneous multi-layered networks is still in its early stages. Multi-layer network describes a network composed of several layers. The layers contain the same nodes but the edges connecting these nodes are different. For example, a gene network may have coexpression links on a first layer and PPI interactions on a second layer. In addition, to depict the relationships between different species of molecules, it is recommended to use heterogeneous networks with a bipartite relationship that describe all possible interaction between two sets of nodes. With more links and more possibilities to fine-tune the transition between layers, more complex network structures would improve the prediction of the random walk [44]. As complex biological networks have several types of molecules and several types of interactions, it becomes essential to use heterogeneous multi-layer networks to describe all the relations of the system. However, tools to build such networks are limited by the number of layers or the number of heterogeneous relations. Therefore, we were encouraged to use monoplex networks for each application. Although monoplex networks highlighted relevant interactions, comparison with heterogeneous multi-layer networks would have been worthwhile.

### Contribution of network-based integration on public datasets

The reanalysis of multi-omics data with new methods can lead to a better understanding of the system. Both cases proposed in this paper are no exception. With the study of HeLa cells during the cell cycle, the authors were able to highlight functional groups, expressed by proteins, which may be over or under-regulated at different phases of the cycle. Combined with the other omics layers, they were able to compare the dynamics of an mRNA, its translation product and the associated protein and thus highlight the regulation mechanism during the cycle. Nevertheless, our methodology, based on network-based multi-omics integration could, in addition to answering the same questions, offer the advantage of going even further.

From a key function (Cell division, Chromosome segmentation, Telomere maintenance) it is possible to identify the molecules involved but also to know the events, interactions, regulation, which have allowed its expression. Moreover, by adding other molecular layers (e.g. metabolites) it is possible to observe the implications of this function at the system level.

The second case examining the response of maize to an aphid attack highlighted the dynamics of certain genes involved in the defense mechanisms, associated with the evolution of known metabolites. Although the initial analyses identified genes and metabolites that are highly variable with the presence of aphids, the authors did not take advantage of all the multi-layer interactions. Indeed, with a very targeted list of metabolites, having roles in defense mechanisms but also in other pathways, it is difficult to identify relevant multi-omic interactions. By adding other layers but also directly connected molecules, we have been able to considerably increase the diversity of the interactions that could be detected and thus increase the chances of identifying other potential targets to better understand maize defense mechanisms against aphids and possibly orient the development of future biological pesticides.

## 5 Conclusion

The widespread use of high-throughput technologies has enabled the profiling of multiple omics layers across multiple time points. This has opened the door to new study designs that can now address questions and identify novel biological mechanisms regulated by different biomolecular layers. However, one of the biggest challenges in this era of multi-omics is the integration and interpretation of the diverse large-scale omics data in a way that provides new and more complete biological insights.

To facilitate the interpretation of longitudinal multi-omics data we have proposed an analytic strategy that consists of multi-omics kinetic clustering and multi-layer network. The use of multi-layer networks can evolve and be enriched by the addition of new layers (for example miRNA, disease, drugs, environmental variables, microbiome) and their respective cross-layered interactions (gene-disease association, host-microbe interaction, miRNA-mRNA regulation) and also intra-layered interactions that could defined different kinds of interactions for the same layer (co-expression, functional relationship, expression delays).

The analysis of two multi-omics longitudinal studies in two previously published datasets demonstrates that new multi-layered interactions can be unravelled. Eventually, and with highly controlled experimental designs, we expect that dynamic networks can be built to model an entire biological system. Therefore, this methodology will open new avenues for exploration and interpretation from multi-omic studies.

## Data Availability Statement

HeLa Cell raw data for case study 1 can be found in [1]. Maize raw data for case study 2 can be found in [42] Processed data and in-house scripts are available in a Github public repository: https://github.com/abodein/netOmics-case-studies

## Supplementary Data

Available as online document.

## Author Contributions

AB performed the analyses and wrote the manuscript. AB, MPSB and KALC revised the manuscript. All authors read and approved the submitted version.

