## supplemental material for "Interpretation of network-based integration from multi-omics longitudinal data"

#### 1 Supplementary Tables

Table 1: HeLa cell cycling study: network statistics of **(A.)** ARACNe gene networks and sub-networks by kinetic clusters. **(B.)** Protein-Protein Interaction sub-networks from BioGRID interaction database with only measured molecules. **(C.)** Same as B. with the addition of BioGRID first degree proteins.

| **(A.)** | Connected Nodes | Isolated Nodes | Edges |
|---|---|---|---|
| Cluster 1 | 223 | 10 | 2338 |
| Cluster 2 | 67 | 14 | 224 |
| Cluster 3 | 25 | 10 | 33 |
| Cluster 4 | 86 | 11 | 884 |
| Entire network | 421 | 25 | 4079 |

| **(B.)** | Connected Nodes | Isolated Nodes | Edges |
|---|---|---|---|
| Cluster 1 | 1,784 | 29 | 19,714 |
| Cluster 2 | 20 | 33 | 15 |
| Cluster 3 | 9 | 41 | 6 |
| Entire network | 1,886 | 30 | 21,679 |

| **(C.)** | Connected Nodes | Edges |
|---|---|---|
| Cluster 1 | 24,300 | 1,177,556 |
| Cluster 2 | 5,327 | 82,438 |
| Cluster 3 | 4,805 | 131,256 |
| Entire network | 24,498 | 1,518,635 |

Table 2: HeLa cell cycling study: Over Representation Analysis results. Significant GO terms (BP, MF, CC) enriched from mRNA, translation products and proteins list set separately. ORA was performed by kinetic clusters and with entire sets of molecules. Additionally, p-values of enriched terms shared between mRNA, translation products and proteins were combined using Fisher's combined probability test. External file : 'table_hela_ora.xlsx'

Table 3: HeLa cell cycling study: Random Walk with restart results on multi-omics network. First sheet ("mechanism"): Top 25 closest nodes from GO terms seeds. Second sheet ("prediction"): Closest GO term node (BP, MF,CC) from unlabelled seeds. Third sheet ("cluster"): Top 10 closest nodes from seeds. Nodes were labelled according to their kinetic clusters. Results were filtered to show only seeds with different clusters from the other nodes. External file : 'table_hela_rwr.xlsx'

Table 4: Dynamic response to maize aphid feeding study: network statistics of ARACNe gene networks and sub-networks by kinetic clusters.

| | Cluster 1 | Cluster 2 | Cluster 3 | Cluster 4 | Entire Network |
|---|---|---|---|---|---|
| Connected Nodes | 180 | 153 | 42 | 64 | 473 |
| Isolated Nodes | 334 | 357 | 48 | 72 | 777 |
| Edges | 204 | 206 | 45 | 61 | 659 |

Table 5: Dynamic response to maize aphid feeding study: Over Representation Analysis results. Significant GO terms (BP, MF, CC) enriched from mRNA, and proteins list set separately. ORA was performed by kinetic clusters and with entire sets of molecules. External file : ‘table_maize_ora.xlsx‘

Table 6: Dynamic response to maize aphid feeding study: Random walk with restart results on multi-omics network. First sheet (”mechanism”): Top 25 closest nodes from GO terms seeds. Second sheet (”prediction”): Closest GO term node (BP, MF,CC) from unlabelled seeds. Third sheet (”cluster”): Top 10 closest nodes from seeds. Nodes were labelled according to their kinetic clusters. Results were filtered to show only seeds with different clusters from the other nodes. External file : ‘table_maize_rwr.xlsx‘

### References
